# Congenital heart disease missense mutations in the TBX5 DNA-binding domain alter thermal stability and DNA-binding affinity

**DOI:** 10.1101/2025.05.16.654568

**Authors:** Alejandro Rivera-Madera, Edwin G. Peña-Martínez, Jean L. Messon-Bird, Diego A. Pomales-Matos, Oswaldo L. Echevarría-Bonilla, Leandro Sanabria-Alberto, Esther A. Peterson-Peguero, José A. Rodríguez-Martínez

**Author notes:** These authors contributed equally to this work (Alejandro Rivera-Madera, Edwin G. Peña-Martínez).

## Abstract

Missense mutations can alter the biochemical properties of proteins, including stability, structure, and function, potentially contributing to the development of multiple human diseases. Mutations in *TBX5*, a transcription factor (TF) necessary for heart development, are among the causes of congenital heart diseases (CHD). However, further research on biophysical and biochemical mechanisms is needed to understand how missense mutations in TFs alter their function in regulating gene expression. In this work, we applied *in vitro* and *in silico* approaches to understand how five missense mutations in the TBX5 T-box DNA-binding domain (I54T, M74V, I101F, R113K, and R237W) impact protein structure, thermal stability, and DNA-binding affinity to known TBX5 cognate binding sites. Differential Scanning Fluorimetry showed that mutants I54T and M74V decreased thermal stability, whereas I101F and R113K had increased stability. Additionally, DNA-binding affinity decreased for all five missense mutants when evaluated *in vitro* for known TBX5 genomic binding sites within regulatory elements of *Nppa* and *Camta1 genes*. Structural modeling of the TBX5 T-box domain predicted altered protein conformation and stability due to the loss or gain of amino acid residue interactions. Together, our findings provide biophysical and biochemical mechanisms that can be further explored to establish causality between TBX5 missense mutations and the development of CHDs.

**Article Summary:** Mutations in genes crucial for heart development, such as *TBX5*, a cardiac transcription factor (TF), have been shown to cause congenital heart diseases. However, the functional consequences of missense mutations in cardiac TF have not been fully explored. In this work, we performed biochemical evaluations and found significant changes in thermal stability and DNA-binding of five TBX5 missense mutants (I54T, M74V, I101F, R113K, and R237W). Additionally, we generated *in silico* predictions of protein structure, stability, and pathogenicity for the five missense mutants. Our approach is scalable to other TF mutants crucial for organ development to further our understanding of congenital disease etiology.

**Graphical Abstract:** 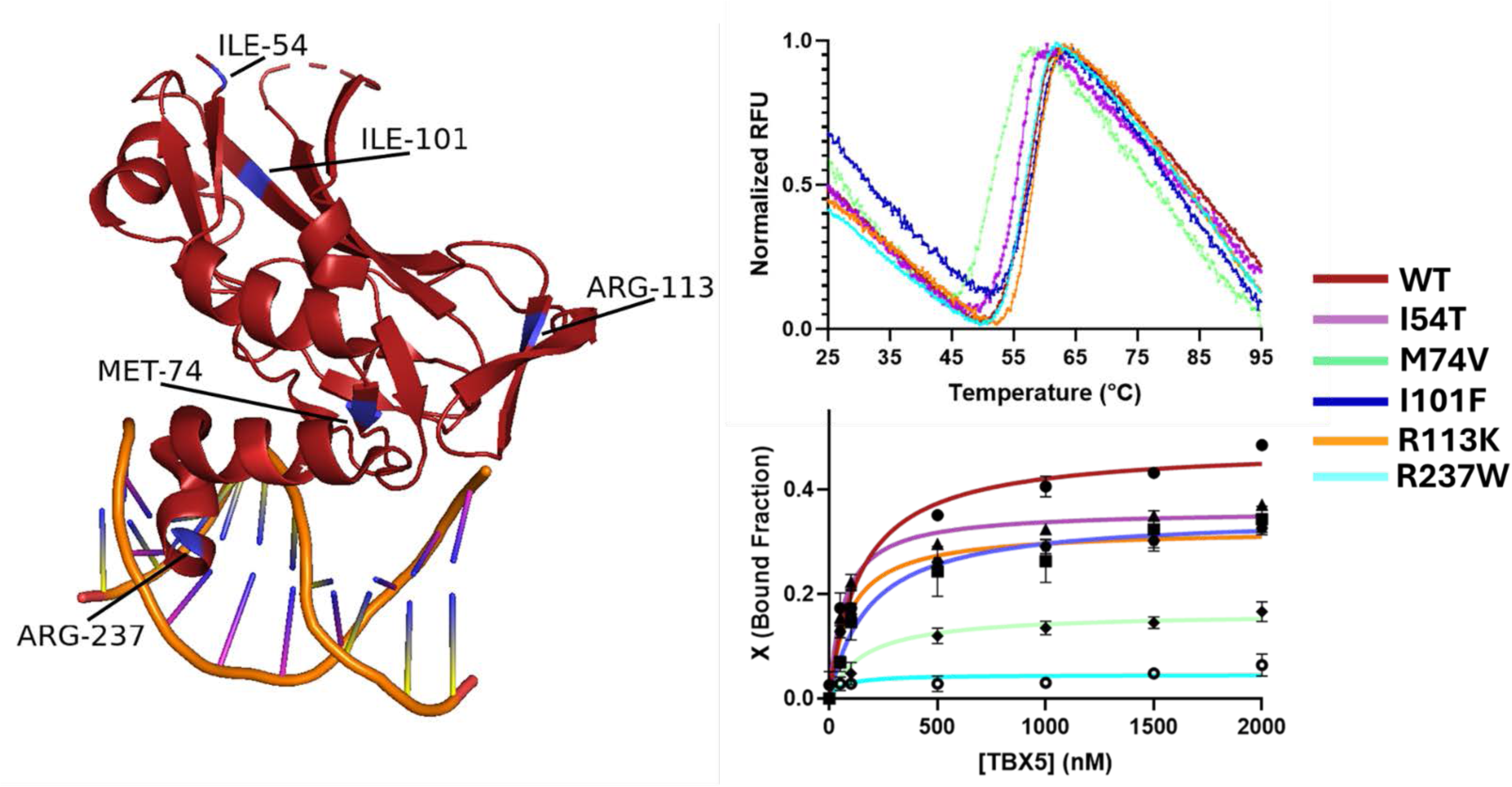

## Introduction

Missense mutations, genetic variants that cause a change in an amino acid, can adversely affect protein structure and function, leading to pathogenic outcomes. [1–3] Missense mutations have been linked to numerous human genetic diseases, such as sickle cell anemia and cystic fibrosis. [4–6] Missense mutations have been shown to alter biochemical mechanisms, such as protein stability, protein folding, protein activity, intermolecular interactions, post-translational modifications, and cellular localization.[7–11] At a cellular level, mechanisms like gene expression have been proven to be altered by missense mutations of regulatory proteins, like transcription factors (TFs). [12–15]

TFs are sequence-specific DNA-binding proteins that regulate the spatial and temporal gene expression. Previous work linking TF mutants to human diseases has reported altered structure, thermal stability, and function of multiple TFs. [12,16–20] For over 30 years, oncogenic missense mutations have been known to alter the stability of p53. [16,17,21] In an *in vivo* system, Kirk *et al.* used *Xenopus laevis* (African clawed frog) embryos model to link clinical TBX20 missense mutations to disruption of developmental processes, such as gastrulation. [19] Although further insight is required to understand how missense mutations can alter TF function and regulatory activity, understanding changes in TF stability and DNA-binding is a step forward in establishing causal mechanisms of many human diseases.

TBX5, also known as T-box transcription factor 5, is an evolutionarily conserved TF involved in heart and limb development. [22–24] The *TBX5* gene is expressed during embryonic development in multiple layers (epicardium, myocardium, and endocardium) and chambers (atrial appendages and left ventricle) of the developing heart. [25,26] TBX5 binds to the consensus sequence 5’-`AGGTGT`-3’, and regulates multiple cardiac genes, such as the *natriuretic peptide precursor A (Nppa)* and *calmodulin binding transcription activator (Camta1)*, to regulate gene expression and promote cell differentiation into cardiac tissue. [27–29] Knockouts of *TBX5* in mice have shown decreased expression of cardiac genes needed for heart development (e.g., *Nppa* and *CX40*) and even developmental arrest or embryo lethality. [30–32] Missense mutations of TBX5 have been associated with multiple types of congenital heart diseases (CHDs), such as Holt-Oram syndrome (HOS), atrial septal defect (ASD), and ventricular septal defect (VSD). [33–37] Missense mutations can alter TBX5 function by impairing molecular interactions with known binding partners (e.g., NKX2-5, GATA4, Mef2C, and SRF) active during heart development. [38–41] Additionally, mutations in the TBX5 DNA-binding domain (DBD) can alter TF-DNA interactions with *cis*-regulatory elements (CRE, e.g., promoters and enhancers) and disrupt the expression of genes crucial for heart development. [13,15,42] For example, one of the first reported TBX5 mutants, R237W, showed decreased binding to the human *ANF* promoter. [22,43] TBX5 mutants have been previously reported to have altered DNA-binding affinity for known binding sites. [22,34,35,44–46]

In this work, we evaluated the thermal stability and DNA-binding activity of five CHD-associated missense mutants in the TBX5 T-box DNA-binding domain (I54T, M74V, I101F, R113K, and R237W; **Figure 1A**). All five CHD-associated missense mutants in the T-box domain of TBX5 were identified from the ClinVar (M74V) [47] and OMIM [48] databases, and most were identified in patients or family trios with HOS (I54T and R237W), ASD (I101F), and VSD(R113K). Recombinant TBX5 T-box missense mutants were expressed and purified through affinity chromatography. Functional evaluation of TBX5 missense mutants was performed through DSF and electrophoretic mobility shift assay (EMSA). We observed changes in TBX5 mutant melting temperature (T_m_) and binding affinity for known TBX5 genomic binding sites. All five variants were computationally modeled to predict structure, amino acid residue interactions, and thermodynamic stability changes (**Figure 1B**). Our results provide the first evidence of altered thermal stability for mutants I54T, I101F, R113K, and DNA-binding affinity for mutants M74V, I101F, and R113K. Furthermore, R237W was used as a control for altered thermal stability and DNA-binding affinity, as was previously described. [22,49] This suggests a role in congenital malformations for CHD-associated missense mutations in TBX5 by disrupting its regulatory function.

**Figure 1:**
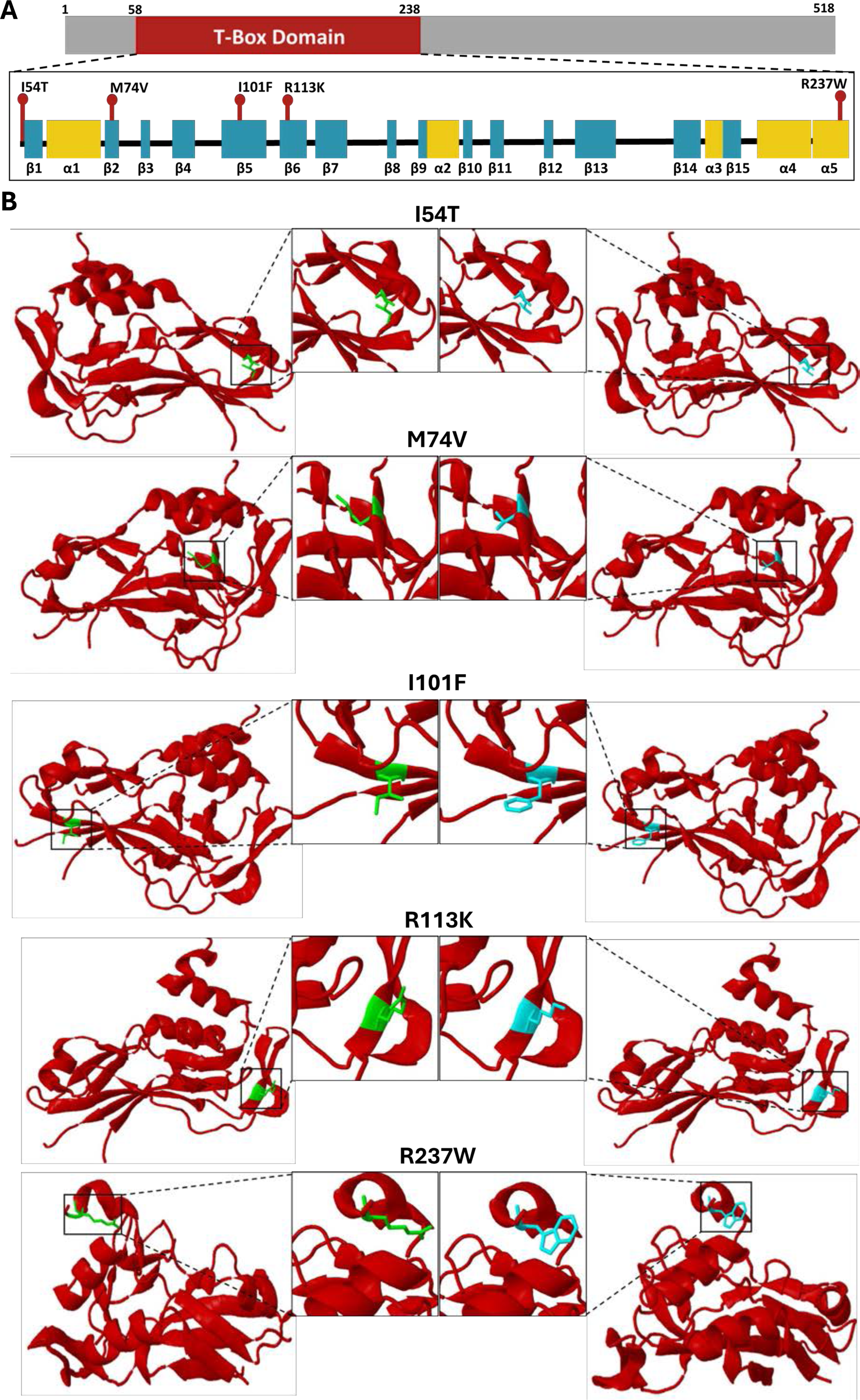
Diagram of TBX5 missense mutants evaluated in this work. **A)** Location of mutations within the TBX5 T-Box Domain. Alpha helices and beta sheets within the T-box domain are represented as yellow and blue rectangles, respectively. **B)** Amino acid side chain substitution in each mutant. Wt residues are on the left in green, and mutant residues are on the right in blue. Modeled with Missense 3D using PDB 2X6V [50].

## Materials & Methods

### Cloning, expression, and purification of TBX5 T-Box Domain

The TBX5 T-box domain plus ten amino acids flanking at each end (Leu48-Ser248) was cloned into the pET-51b(+) bacterial overexpression vector using Gibson Assembly (New England Biolabs, #E5510S). pET51b(+) adds an N-terminal Strep-Tag-II and C-terminal 10X His-Tag. Five missense mutations were generated using the Quick-change II Site-Directed Mutagenesis kit (Agilent Technologies, #200523). All plasmids were verified through long-read whole-plasmid sequencing (Plasmidsaurus Inc.) and transformed into BL21 DE3 *E. coli* (New England Biolabs #C2530H or Millipore Sigma #70956-4). First, 50 mL Luria Broth (Sigma-Aldrich, #L3022) cultures were grown for 16 hours at 37°C, shaking at 230 rpm, followed by inoculation of 500 mL of Terrific Broth (Sigma-Aldrich, #T0918) with 10 mL of the overnight culture and grown at 37°C. Once the culture reached OD_600_ between 0.5 and 0.8, protein expression was induced by adding 1 mM IPTG and culturing at 18°C while shaking. After 20 hours, the culture was centrifuged at 2800 x g for 5 minutes at 4°C, and the pellet was frozen at -80°C overnight.

The pellet was resuspended with 40 mL of column buffer, and 4 mL (10% of volume) of 5 M NaCl were added. The sample was then sonicated for four cycles at 40 % amplitude for 30 seconds (QSONICA, Part No. Q125), centrifuged (2800 x g, 30 min, 4°C), and the supernatant was incubated with 2 mL of Ni-NTA resin (Qiagen, # 30210) (1 hour, 4°C, orbital shaking). The column was equilibrated with 20 mL of column buffer (500 mM NaCl, 20 mM Tris-HCl, pH 8.0, 0.2 % Tween-20, 30 mM imidazole, and EDTA-free protease inhibitor (ThermoScientific # A32965). The supernatant was then passed through the column twice, followed by three washes with increasing concentrations of imidazole (30 mM, 50 mM & 100 mM) in column buffer. The protein was eluted six times with 1.8 mL of elution buffer (500 mM NaCl, 20 mM Tris-HCl, pH 8.0, 0.2 % Tween-20, 500 mM imidazole). Buffer exchange to binding buffer (50 mM NaCl, 10 mM Tris-HCl, pH 8.0, and 10 % glycerol) was performed using Amicon® Ultra Centrifugal Filter, 3 kDa concentrators (Millipore Sigma, #UFC5003).

Finally, the purification was evaluated with SDS-PAGE and Western Blot using Mini-PROTEAN TGX Precast Protein Gels® (Bio-Rad, #4561086) as seen in **Supplementary Figure 1**. Samples were prepared with 4X Laemmli Sample Buffer (Biorad #1610747) containing β-mercaptoethanol (BME) for a total volume of 20 μL (5 μL 4X loading buffer + 15 μL sample), heated at 95 °C for 5 minutes, and resolved for 1 hour at 120 V at room temperature. The gel was stained using Coomassie Brilliant Blue R stain (Sigma-Aldrich, #B0149). For the Western Blot analysis, contents from the SDS-PAGE gel were transferred to a PVDF membrane using the Bio-Rad Turbo Transfer System protocol in a Trans-Blot® Turbo™ for 3 min at 25 V. Membranes were blocked using 5 % milk in 1X TBST buffer for 1 hour in orbital shaking and incubated overnight with 1:10,000 dilution of horse radish peroxidase-conjugated Anti-His mouse monoclonal antibody (Novus Biologicals, AD1.1.10). The SDS-PAGE and Western Blot Analysis results were imaged using Azure Sapphire Biomolecular Imager (Azure Biosystems).

### Differential Scanning Fluorimetry

Differential Scanning Fluorimetry (DSF) [51] was used to evaluate the thermal stability of the TBX5 T-box domain and its missense mutants with the Applied Biosystems™ QuantStudio™ 3 Real-Time PCR System, 96-well, 0.2 mL (#A28137) using MicroAmp Optical 8-Tube strips (Applied Biosystems #4316567 & #432302). Samples were heated from 25°C to 95°C with a ramp of 0.05 °C/s and measured with a 470 ± 15 nm / 520 ± 15 nm excitation/emission filter. Protein stocks of 10 µM in binding buffer (50 mM NaCl, 10 mM Tris-HCl, pH 8.0, and 10 % glycerol) were diluted to 5 µM in a 20 µL reaction with 5X SYPRO Orange Dye (Invitrogen™ S6650), and the volume was completed with binding buffer. Melting curve raw data from the DSF melting assays was exported from the equipment and analyzed using DSFworld. [52] Normalized raw data was plotted against the temperature (°C) using GraphPad Prism 10 to obtain melting curves for each wt-mutant pair. T_m_ values and graphs represent the average of 6-9 replicates per protein. Changes in melting point (ΔT_m_) were obtained by subtracting the wt T_m_ from the mutant T_m_ (**Supplementary Figure 3**). In addition, statistical significance was determined through an unpaired parametric t-test of each mutant compared to the wild type.

### Electrophoretic Mobility Shift Assays

DNA-binding activity was assessed through Electrophoretic Mobility Shift Assays (EMSA) with two known binding sites for TBX5. Assays were performed with 40 bp sequences derived from the *natriuretic peptide precursor A* (*Nppa*) and the *calmodulin-binding transcription activator* (*Camta1)* promoters, with an additional 20 bp constant region to add IRDye® 700 fluorophore (Integrated DNA Technologies) through a primer extension reaction (Available in Supplementary Table #2). Binding reactions of 20 µL in binding buffer (50 mM NaCl, 10 mM Tris-HCl, pH 8.0, and 10 % glycerol) were prepared with seven protein concentration points (0, 50, 100, 500, 1000, 1500, and 2000 nM). Acrylamide gels went through a pre-run at 85 V for 15 min before loading samples at 35 V. Samples were incubated for 30 min at 30°C and 30 min at room temperature before resolution in a 6% native polyacrylamide gel for 1.5 hours at 75 V in 0.5X TBE (89 mM Tris HCl, 89 mM boric acid, 2 mM EDTA, pH 8.4). The results were imaged using the Azure Sapphire Biomolecular Imager with an excitation/emission of 658 nm/710 nm (Azure Biosystems) and analyzed as previously described. [53,54] Fold change in the bound fraction was calculated by dividing the maximal binding fraction (B_max_) value of the wt TBX5 T-box domain by the B_max_ value of the corresponding mutant.

### *In silico* prediction of missense mutant’s thermodynamical stability and structure

Predictions of changes in thermodynamic stability were performed for the missense mutants compared to the wt TBX5 using web-based programs that calculate changes in Gibson’s free energy (ΔΔG) and/or protein structure. We predicted thermodynamic stability using DynaMut2 [55], RaSP [56], PremPS [57], iMutant 2.0 [58], and mCSM [59]. Changes in amino acid interactions and structural damage predictions were performed with DynaMut2 [55] and Missense3D [50], respectively (**Figure 1B and 4A**). All predictions were performed using PDB 2X6V and chain A, which represents the protein in bound conformation to the DNA.

In addition, we modeled all six proteins with ColabFold v1.5.5: AlphaFold2 using MMseqs2 and established alignments between wt and each mutant for the structure ranked first using TM-Align (https://seq2fun.dcmb.med.umich.edu//TM-align/). [60] This tool helped us to evaluate the similarity of the folding position of each residue by calculating the TM-Score, which varies in a range from 0-1, where 1 indicates a perfect similarity between the compared structures. All structure predictions were done without both affinity tags. Pathogenicity likelihood was predicted using MutPred2 [61] using the 2X6V TBX5 structure **(Supplementary Table 3)**.

**Reagent Table**:

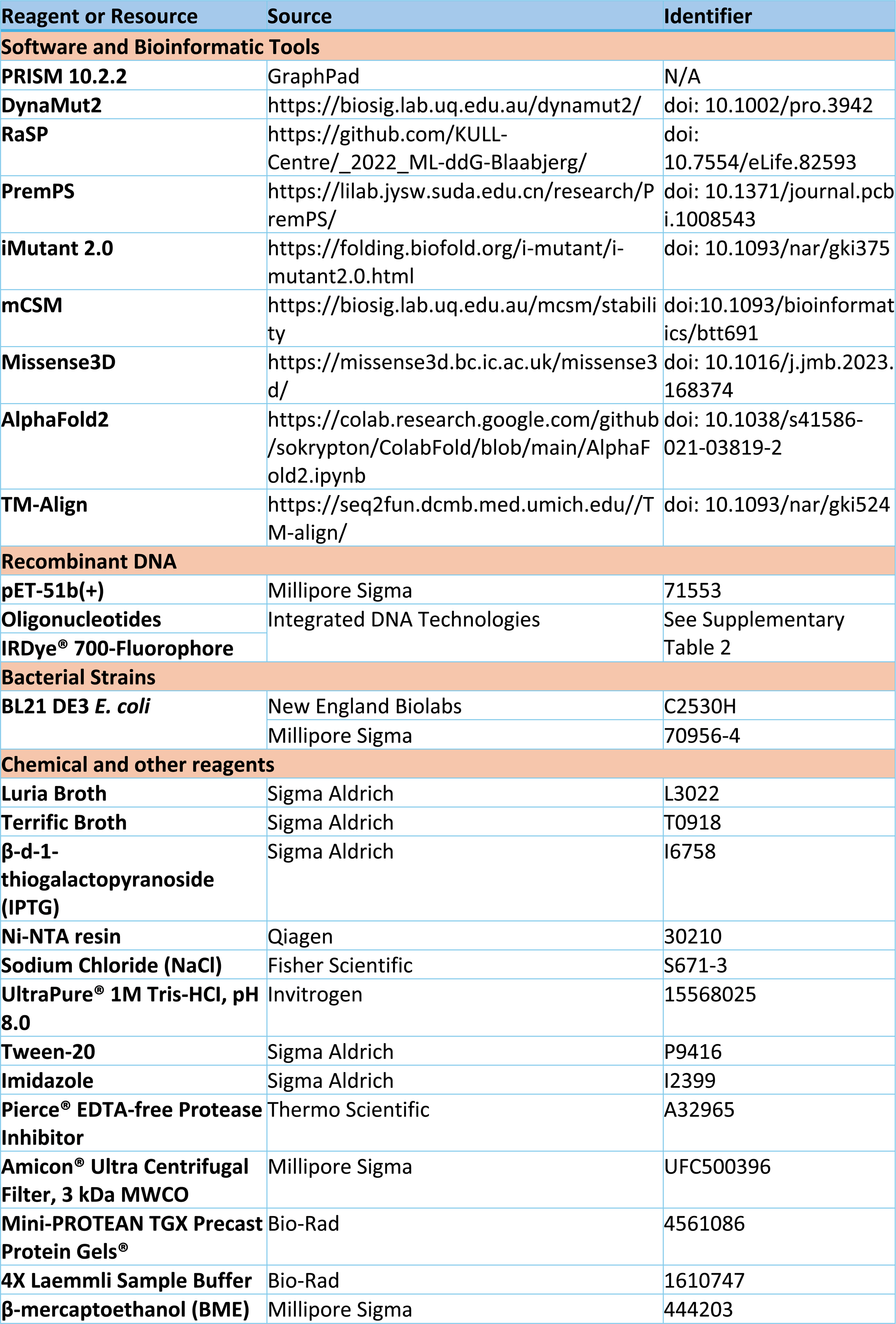
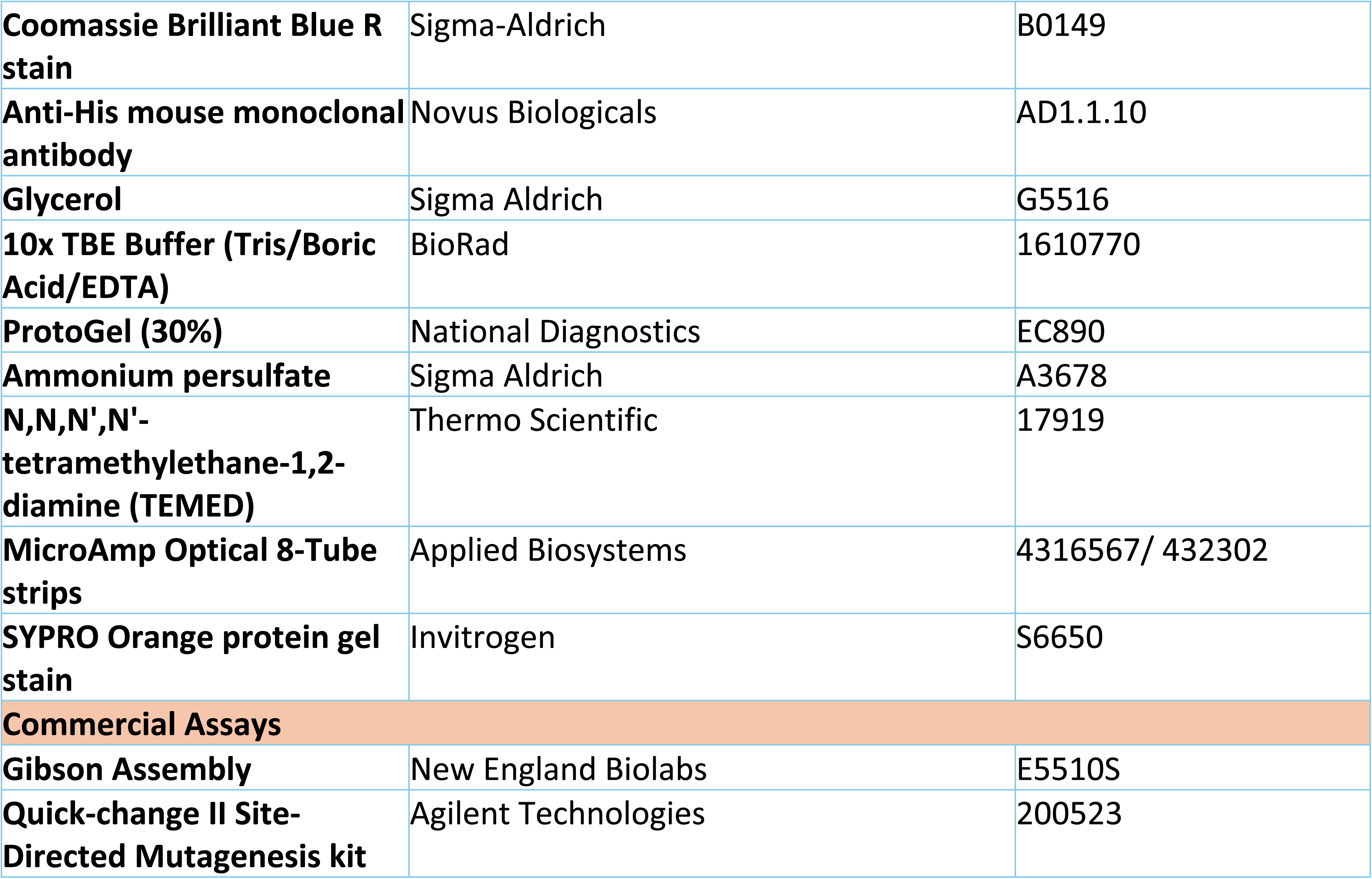

## Results

### Missense mutations in TBX5 T-Box Domain alter thermal stability

We evaluated the *in vitro* thermal stability of the wt TBX5 T-box domain and five missense mutants (I54T, M74V, I101F, R113K, and R237W) with differential scanning fluorimetry (DSF). In DSF, protein thermal stability is determined by measuring the fluorescence of SYPRO Orange during protein thermal denaturation. Changes in thermal stability were determined by comparing the melting temperatures (T_m_) between wt TBX5 and mutants. Four of the evaluated mutants had statistically significant changes in melting temperatures (T_m_) (**Figure 2**). Wt TBX5 T-box domain has a T_m_ of 57.9 ± 0.2. Mutants I54T and M74V showed a decrease in thermal stability with T_m_ values of 55.9 °C ± 1.1 (Δ T_m_ = -2.0 °C) and 53.2 °C ± 0.2 (Δ T_m_ = -4.7 °C), respectively. In contrast, mutants I101F and R113K showed an increase in thermal stability with T_m_ values of 58.9 °C ± 0.3 (Δ T_m_ = 1.0 °C) and 59.1 °C ± 0.5 (Δ T_m_ = 1.2 °C), respectively. No significant changes in thermal stability were observed for R237W, as evaluated through a t-test (**Figure 2B**). Changes in thermal stability for each mutant are summarized in **Table 1**.

**Figure 2:**
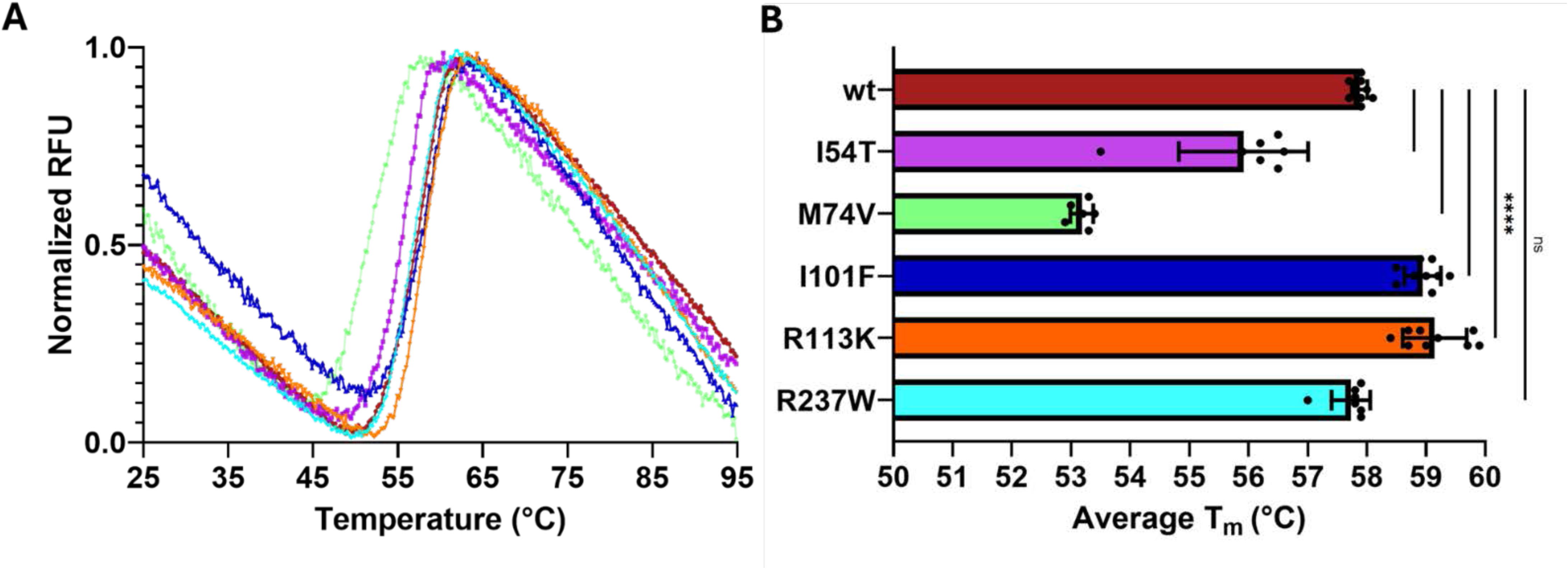
Evaluation of mutant TBX5 T-box domain thermal stability by differential scanning fluorimetry. **A**) Melting curves were generated for the wt T-box domain and each missense mutant. Melting curves normalization and T_m_ analysis were performed using DSFworld. RFU: relative fluorescent units **B**) T_m_ quantification and statistical analysis through unpaired t-test (**** = p-value < 0.0001). Color scheme: Wt T-box domain (red), I54T (purple), M74V (green), I101F (blue), R113K (orange), and R237W (cyan).

**Table 1:**
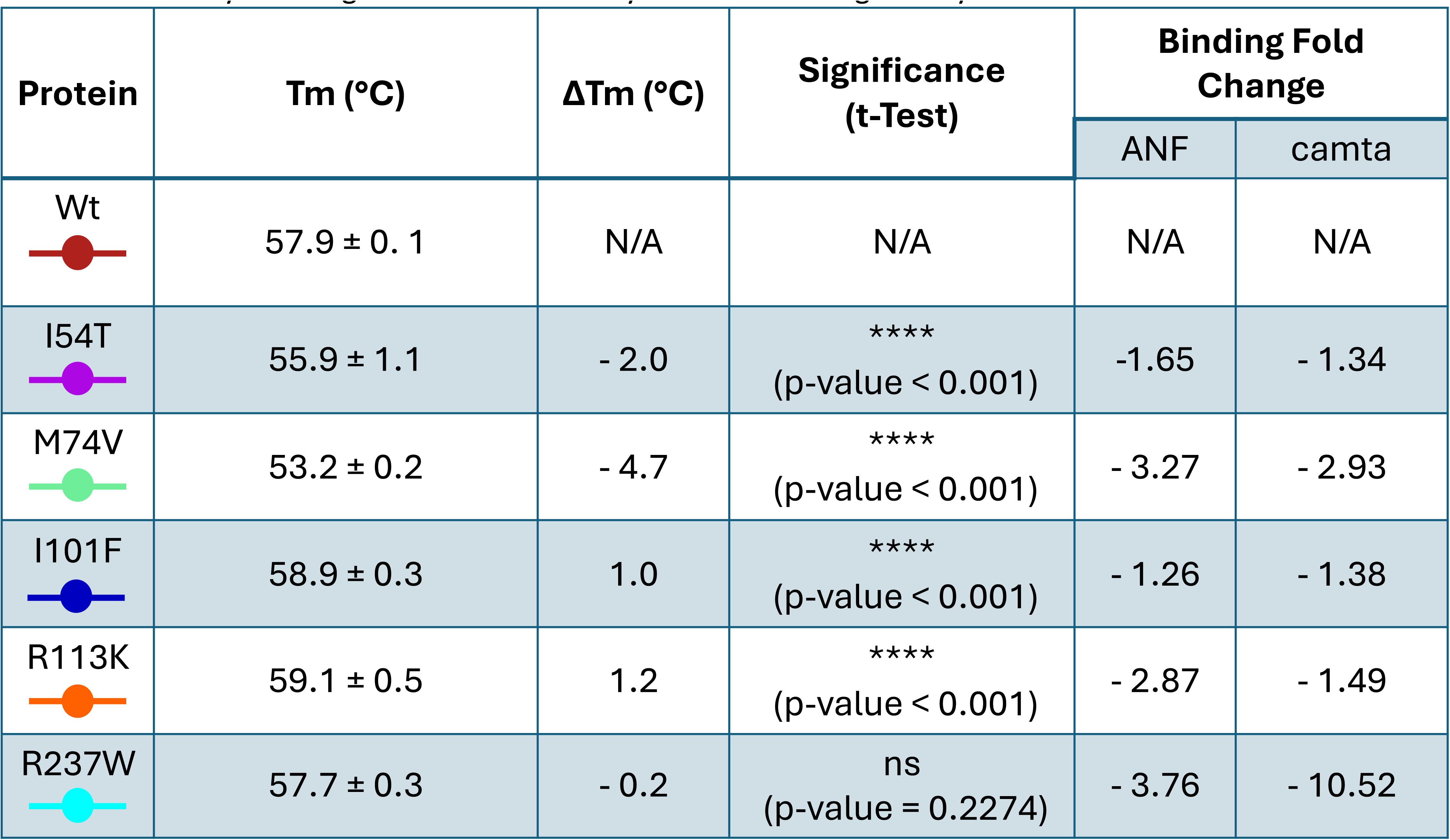
Summary of changes in thermal stability and DNA-binding affinity validated *in vitro*.

| Protein | Tm (°C) | ΔTm (°C) | Significance<br>(t-Test) | Binding Fold Change |  |
| --- | --- | --- | --- | --- | --- |
|  |  |  |  | ANF | camta |
| Wt | 57.9 ± 0.1 | N/A | N/A | N/A | N/A |
| I54T | 55.9 ± 1.1 | - 2.0 | ****<br>(p-value < 0.001) | -1.65 | - 1.34 |
| M74V | 53.2 ± 0.2 | - 4.7 | ****<br>(p-value < 0.001) | - 3.27 | - 2.93 |
| I101F | 58.9 ± 0.3 | 1.0 | ****<br>(p-value < 0.001) | - 1.26 | - 1.38 |
| R113K | 59.1 ± 0.5 | 1.2 | ****<br>(p-value < 0.001) | - 2.87 | - 1.49 |
| R237W | 57.7 ± 0.3 | - 0.2 | ns<br>(p-value = 0.2274) | - 3.76 | - 10.52 |

### TBX5 missense mutants decrease binding affinity for known binding sites

To further determine the biochemical implications of these missense mutations, we analyzed changes in binding affinity for known TBX5 genomic binding sites. TBX5 T-box domain DNA-binding affinity was evaluated *in vitro* through electrophoretic mobility shift assay (EMSA) using fluorescently labeled oligonucleotides containing 40 bp derived from the *natriuretic peptide precursor A* (*Nppa*) and *calmodulin binding transcription activator* (*Camta1*) promoter. Binding curves were constructed using seven protein concentrations (0, 50, 100, 500, 1000, 1500, and 2000 nM) and averaging three experimental replicates.

All five missense mutants showed a decrease in DNA-binding affinity to both the *Nppa* (**Figure 3A**) and the *Camta1* promoters (**Figure 3B**) compared to the wt TBX5 T-box domain. Mutant R237W had the greatest impact on DNA-binding, with a 3.76- and 10.52-fold change for known binding sites within the *Nppa* and *Camta1* promoters, respectively. This was followed by M74V with a 3.27- and 2.93-fold change decrease for *Nppa* and *Camta1*. Mutant I54T had a 1.65- and 1.34-fold decrease for *Nppa* and *Camta1*, respectively. Mutant R113K had a 2.87- and 1.49-fold change decrease for *Nppa* and *Camta1*. Finally, mutant I101F had the lowest impact on DNA binding with a 1.26- and 1.38-fold change decrease for *Nppa* and *Camta1*, respectively. Fold changes in binding for each mutant are summarized in **Table 1**.

**Figure 3:**
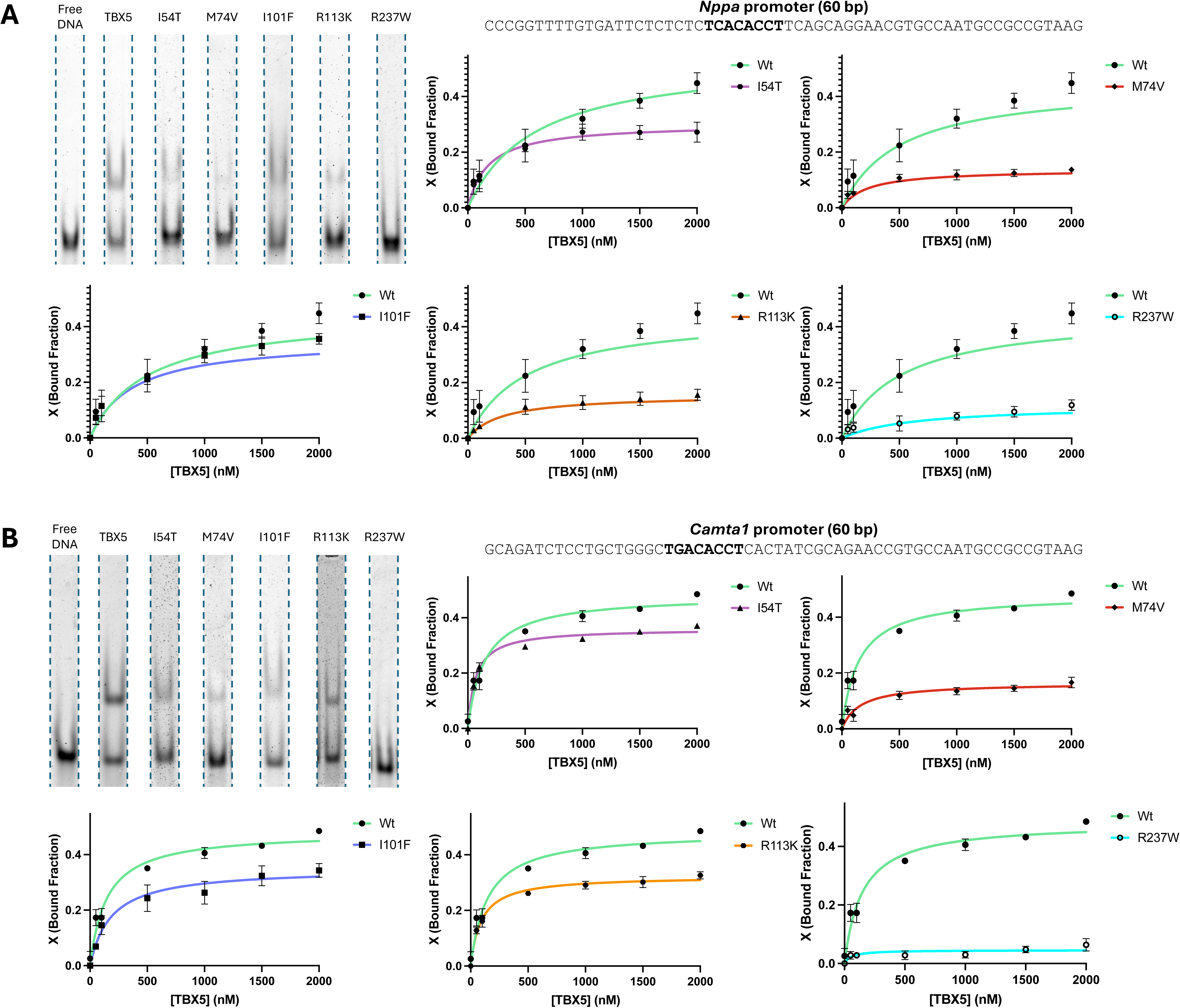
Evaluation of *in vitro* DNA-binding affinity on TBX5 missense mutants for known binding sites. Binding assays for **A**) mouse *natriuretic peptide precursor A* (*Nppa*) and **B**) mouse *calmodulin binding transcription activator* (*Camta1*) promoters. The figure shows EMSA gel shift bands at 2000 nM (upper left) and binding curves for each T-box mutant. Gel images were derived from **Supplementary Figure 4**. Color scheme: Wt T-box domain (red), I54T (purple), M74V (green), I101F (blue), R113K (orange), and R237W (cyan). Sequences used for fluorescent probes are at the top of each figure with the wt TBX5 binding motif in bold.

**Figure 4:**
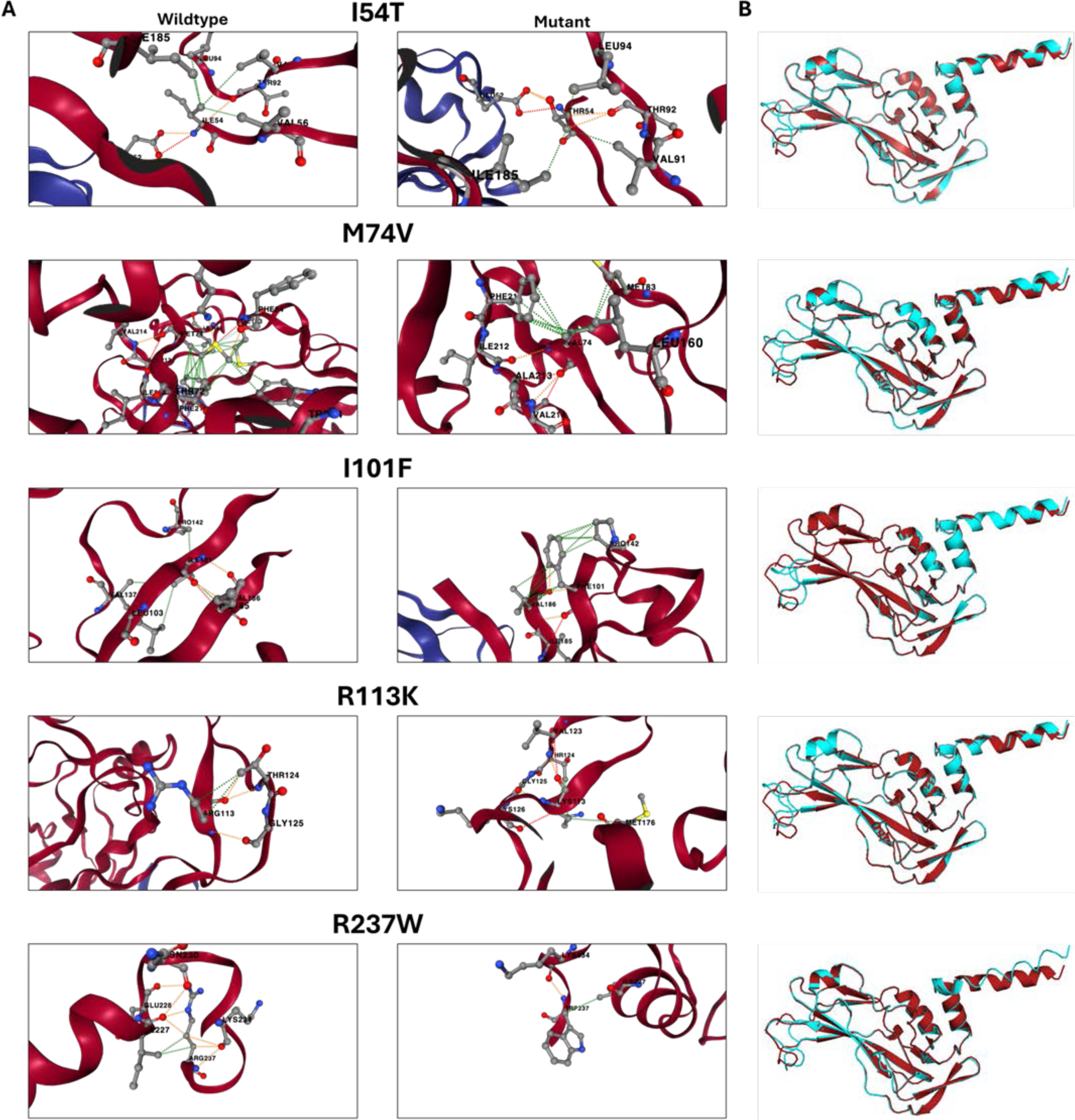
*In silico* modeling of amino acid interaction of the TBX5 T-box domain. **A**) Interactions between amino acids of wt T-box domain (left) with its mutant counterpart (right). Images are zoomed in and centered on the amino acid substitution. Specific interactions are illustrated as follows: Hydrophobic interactions (green), Hydrogen bonds (red), polar bonds (orange), ionic bonds (yellow), and Van der Walls (light blue). **B**) Structural similarities between the T-box domain and missense mutants. Structures were predicted using Alphafold2 and superimposed with TMAlign: wt T-box domain (red), and mutant (blue).

### TBX5 missense mutations are predicted to decrease thermodynamic stability and alter structural features

We performed predictions of changes in thermodynamic stability (ΔΔG), structural damage, changes in amino acid interactions, and 3D structure using tools with web user interfaces (**Table 2**). All mutations were predicted to decrease the thermodynamic stability relative to the wt TBX5 T-box domain. Mutant M74V was the only mutation predicted to have structural damage, with the loss of a buried hydrogen bond donated from Lys226 and a cavity alteration, as predicted with Missense3D. Additionally, all TBX5 missense mutants were predicted to be pathogenic through the gain or loss of biochemical features (e.g., loss of post-translational modifications, tertiary structures, etc.) and protein function (e.g., DNA binding, metal binding, allosteric/catalytic sites, etc.) (**Supplementary Table 3**). Mutant M74V had the highest pathogenicity score (0.92/1.00), with predicted altered DNA/metal binding and PTMs sites (e.g., methylation site in lysine 78). Conversely, I54T was the least pathogenic (0.72/1.00), which predicted altered metal binding and stability. Among the remaining mutants, I101F (0.79/1.00) had a predicted loss of acetylation and methylation at lysine 99 and altered metal binding capacity. Mutants R113K (0.80/1.00) and R237W (0.87/1.00) had new acetylation and methylation sites, and altered DNA-binding activity. Our *in vitro* findings support the altered stability predicted for I54T and DNA-binding for M74V, R113K, and R237W. Protein interactions are mediated by contacts with specific amino acid residues, such as TF binding to DNA or enzymatic recognition for post-translational modifications. As expected, protein function and biochemical properties can change by altering specific amino acid interactions with other molecules.

**Table 2:**
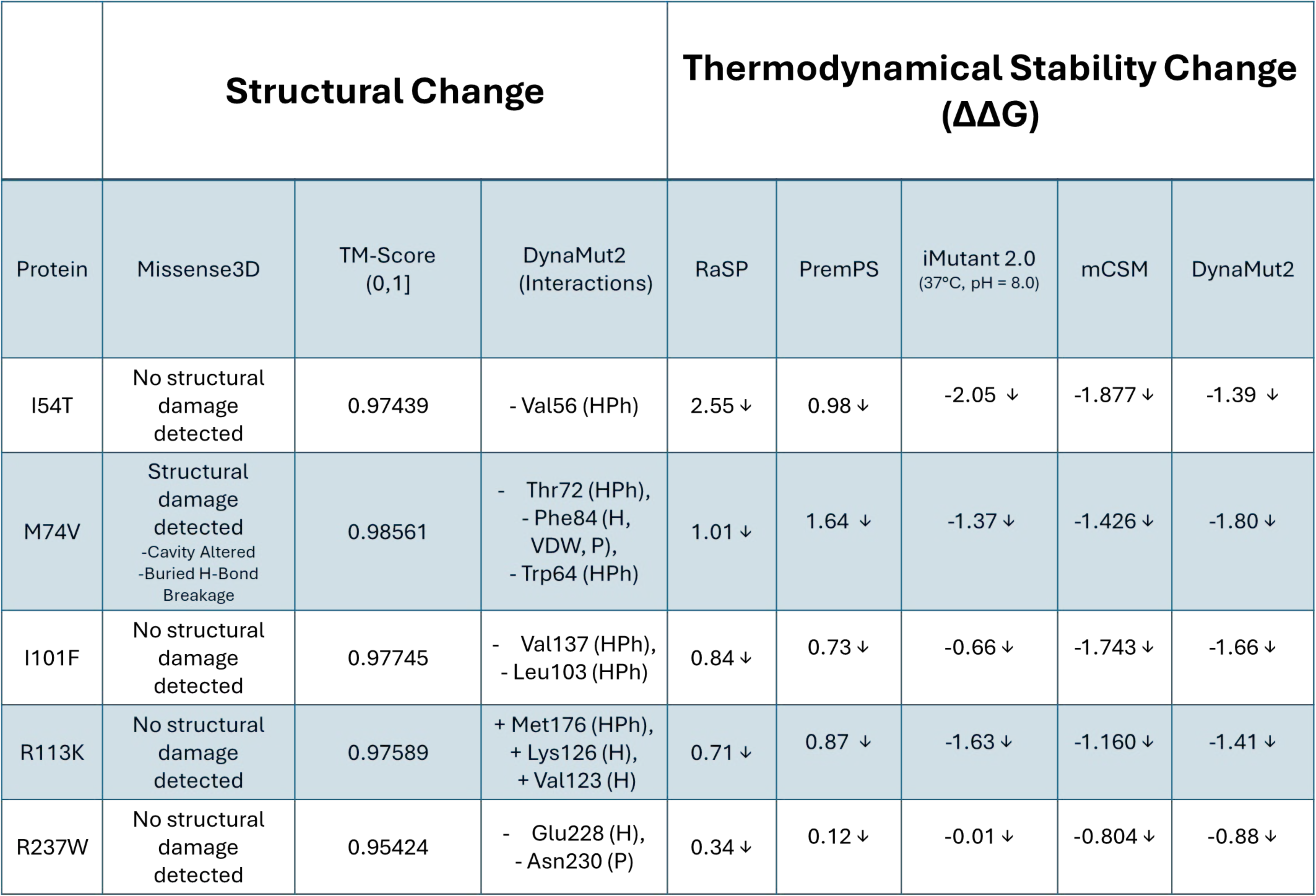
*In silico* prediction on structural and thermodynamic stability changes for T-box domain mutants. New amino acid interactions are marked with a plus (+), while broken interactions are marked with a minus (-). Specific amino acid interactions are described as follows: Hydrophobic (HPh), Hydrogen (H), Van der Waals (VDW), and Polar (P). Mutants with a predicted decrease in thermodynamic stability are marked with a downward arrow (↓) next to the computational score.

To further understand these predictions, amino acid interactions in the TBX5 T-box domain were visualized using DynaMut2 and compared to all five missense mutants (**Figure 4A**). DynaMut2 is a machine learning model that predicts the effects of point mutations on protein stability and dynamics through protein folding and structure analysis. All mutants resulted in at least one broken or new interaction between neighboring amino acid residues. When compared to the wt TBX5 T-box domain, mutant I54T lost a hydrophobic interaction with V56. Mutant M74V lost two hydrophobic interactions with residues T72 and W64, and three interactions with residue F84 (Hydrogen bond, Van Der Waals & Polar). Mutant I101F lost two hydrophobic interactions with residues V137 and L103. Mutant R113K was the only one predicted to create new interactions by forming two new hydrogen bonds with residues K126 and V123, and a hydrophobic interaction with M176. Mutant R237W lost a hydrogen bond with residue E228 and a polar interaction with residue N230. Finally, we predicted structures for the wt and mutant proteins using Alphafold2 and compared them using TmAlign (**Figure 4B** and **Supplementary Figure 2**). TMScore values (0,1] were all >0.95, which indicates a high similarity in the folded structure between all the mutant structures and the wt.

## Discussion

Rare coding variants in genes that are essential for heart development, such as *TBX5*, *NKX2-5*, and *GATA4*, contribute to the etiology of congenital heart diseases (CHDs). [23,45,62–64] Hence, understanding the molecular mechanisms of mutations in proteins needed for proper heart development, such as cardiac TFs, is key to understanding CHD etiology. In this work, we aimed to understand how the missense mutants of the cardiac TF TBX5 can alter its thermodynamic stability, DNA-binding properties, and structural properties. Towards this goal, we focus our work on five TBX5 missense mutations (I54T, M74V, I101F, R113K, and R237W) in the DNA-binding domain present in CHD patients from ClinVaR [47] (**Figure 1A-B**). Towards this goal, we applied a series of computational tools to perform *in silico* predictions on structural changes [46,47] and thermodynamic stability [55–59] for all five TBX5 missense mutants and validated them *in vitro* (**Tables 1 and 2**).

First, we evaluated how these predicted structural changes may alter TBX5 stability and function, we cloned, expressed, and purified wt and mutant TBX5 T-box domains (**Supplementary Figure 1**). We performed thermal stability assays on the wt T-box domain and all five missense mutants to evaluate changes in their T_m_ (**Figure 2**). Two missense mutations (I54T and M74V) decreased thermal stability, while two (I101F and R113K) resulted in an increase after *in vitro* evaluation through DSF. M74V had the greatest decrease in thermal stability (ΔT_m_ = -4.7 °C), which was expected since it is a large hydrophobic residue buried in the hydrophobic core and there are five predicted disrupted interactions, leading to structural damage as predicted by Missense3D. Conversely, R113K had the highest gain in thermal stability (ΔT_m_ = 1.2 °C) while being the only mutant with novel predicted residue interactions. In these cases, with opposite outcomes, M74V and R113K had a respective loss and gain of hydrophobic interactions and hydrogen bonds, which greatly contribute to protein folding and structural stability, respectively. [65–67] Similarly, mutants I54T and R237W resulted in a decrease in thermal stability (ΔT_m_ = -2.0 and -0.2 °C, respectively), while I101F increased (ΔT_m_ = 1.0°C). Overall, three of the thermodynamic stability predictions were consistent (I54T, M74V, and R237W), which decreased just as all the computational tools predicted. The findings and data available from this work (available in **Supplementary File 1**) can be used to train or improve existing models to cover some of the current limitations in predicting protein stability at a single amino acid resolution.

To further understand how missense mutations may alter TBX5 function, we next evaluated changes in DNA-binding affinity to known TBX5 genomic binding sites. Through *in vitro* binding assays, we observed a decrease in affinity to known TBX5 binding sites within the *Nppa* and *Camta1* promoters, two genes involved in heart development (**Figure 3, Supplementary Figure 4**). Mutant M74V had the greatest decrease in DNA-binding affinity, with a 3.27- and 2.93-fold decrease for *Nppa* and *Camta1*, respectively. This decrease may be derived from the significant impact on M74V thermal stability as well as the predicted structural damage, limiting the formation of interactions needed for TF-DNA binding. However, the greatest decrease in binding affinity for both *Nppa* and *Camta1* probes came from mutant R237W. Substitution of the positively charged arginine residue to the non-polar bulky tryptophan can hinder the formation of ionic interactions with the negative charges of the DNA phosphate backbone. [68–70] Similarly, mutant R113K also had an arginine residue substitution and considerably decreased affinity for both the *Nppa* and *Camta1* sequences. Although the substitution was for a lysine, another positively charged residue, previous work has shown that arginine-rich domains are important for the binding of some TFs. [71–74] Mutants I54T and I101F decreased the DNA-binding affinity of the T-Box domain while showing decreased and increased thermal stabilities, respectively. These findings are consistent with previous work done in TBX5 missense mutants (reduced thermal stability of M74V and DNA-binding for I54T and R237W) and can complement past research on protein-protein interactions, gene expression, and cellular localization. [22,43,49,75] Through this approach, we have demonstrated that missense mutations within the T-Box domain of TBX5 can impact TF-DNA binding affinity. Not only can mutant TFs lose affinity for known binding sites, but they also open the possibility of having novel binding sites in the human genome and dysregulating genes outside their regulatory network. [15,76,77]

All five missense mutant structures are predicted to have changes in amino acid residue interactions within the T-box domain (**Figure 4**). Four mutants (I54T, M74V, I101F, and R237W) lost residue interactions important for protein secondary structures formation and maintenance (e.g., hydrogen bonds, hydrophobic interactions, etc.), whereas one mutant (R113K) gained interactions. Additionally, computational modeling of biochemical features such as altered stability (I54T), metal/DNA binding (I54T, M74V, I101F, R113K, and R237W), structural features/folding (M74V, I101F, R113K, and R237W), and post-translational modification (M74V, I101F, R113K, and R237W). As such, all five missense mutations were predicted as likely pathogenic mutants (**Supplementary Table 3**).

To summarize, we evaluated the impact of TBX5 missense mutations on protein structure, thermal stability, and DNA-binding affinity through a combined *in silico* and *in vitro* approach. Computational T-box domain structure modeling revealed insights into amino acid residue interactions with potential implications for protein folding and stability. Through thermal stability and DNA-binding assays, we observed changes in the TBX5 T-box domain properties crucial to maintaining biological function. Together, our findings provide biophysical and biochemical mechanisms that can be further explored to establish causality between TBX5 missense mutations and the development of CHDs. We found no direct correlation between altered thermal stability and DNA-binding activity. Only three out of five mutants (I54T, M74V, and R237W) showed proportional changes between thermal stability and binding. Furthermore, we provide novel findings on thermal stability (I54T, I101F, R113K, and R237W) and DNA binding (M74V and R113K). Our comprehensive approach is scalable to other mutations in TFs crucial for organ development, furthering research toward diagnosis and treatments for multiple types of birth defects.

## Acknowledgment

We thank Jessica M. Rodríguez-Ríos for providing the fluorescent probes for the EMSA and Emanuel A. Carrasquillo-Dones for cloning the TBX5 wt construct. This project was supported by NIH-SC1GM127231, NSF [1736026], University of Puerto Rico Rio Piedras Institutional Funds (FIPI), Puerto Rico Science, Technology, and Research Trust, and NIH Institutional Development Award (IDeA) INBRE [P20GM103475]. ARM was funded by NSF REU: PR-CLIMB Program (2050493), NSF EPSCoR Data Outcomes Collection System (EDOCS) 1736026, and NIH 1T34GM145404. EGPM was funded by the NSF BioXFEL Fellowship (STC-1231306). EGMP and DAPM were funded by the NIH RISE Fellowship (5R25GM061151–20). JLMB and ARM were funded by NSF PR-LSAMP fellowship (HRD-2008186). DAPM was funded by NSF [IQ BIOREU 1852259]. OLEB was funded by NSF PR-LSAMP Bridge to the Doctorate Program (2306079). LSA was funded by NIH ID-GENE Fellowship (1R25HG012702–01). The graphical abstract was made using Biorender.

## Author Contribution

ARM, EGPM, and JARM conceived and designed the study. ARM, EGPM, JLMB, DAPM, OLEB, and LSA performed the experiments. ARM, EGPM, and OLEB were involved in recombinant protein expression and purification. ARM and JLMB were involved in the thermal stability assays. ARM, EGPM, DAPM, and LSA performed EMSA. ARM and EGPM were involved in computational prediction, structural modeling, and statistical analysis. ARM and EGPM wrote the original manuscript. EAPP and JARM supervised the work and secured funding. All authors read and approved the manuscript.

---

**Supplementary Figure 1:**
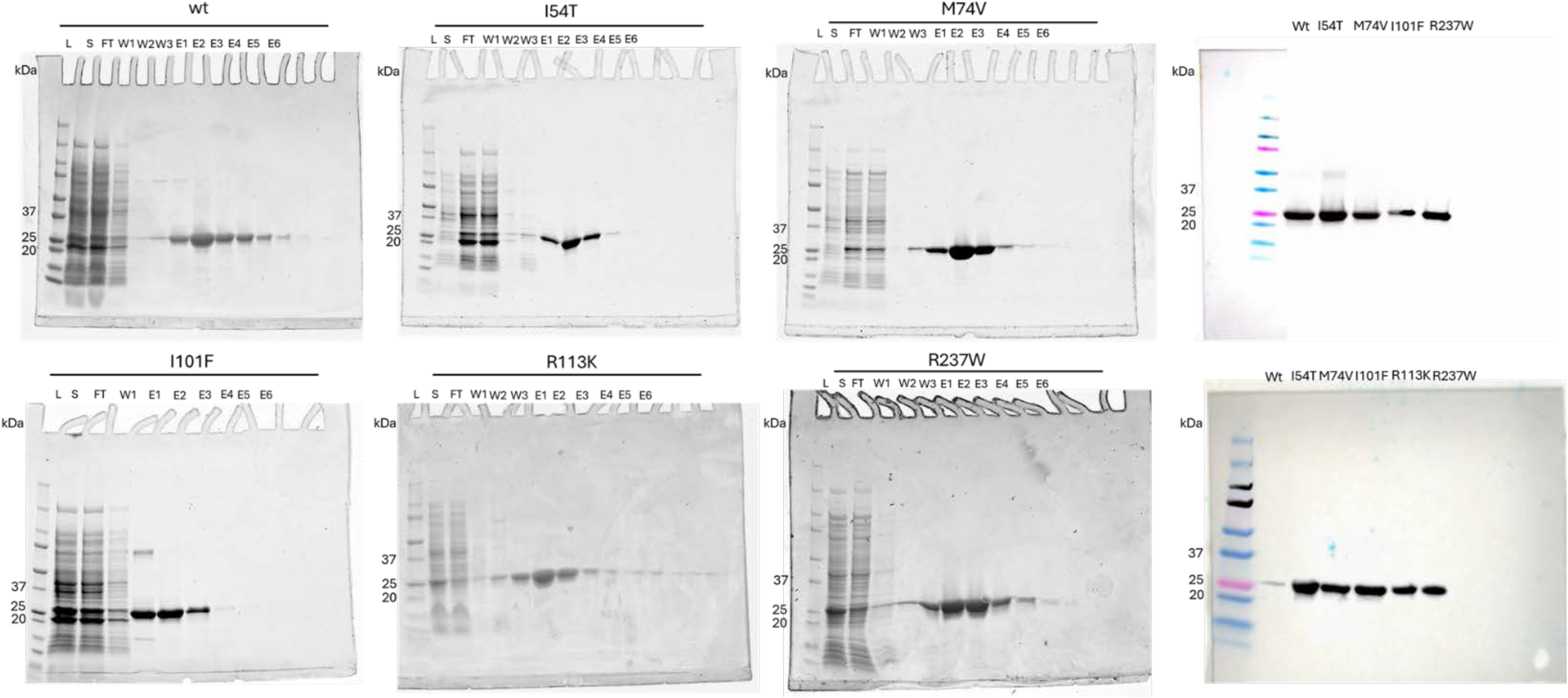
T-box domain and mutant expression, purification, and western blot. Ladder (L), Supernatant (S), Flowthrough (FT), Wash (W), Elutes (E).

**Supplementary Figure 2:**
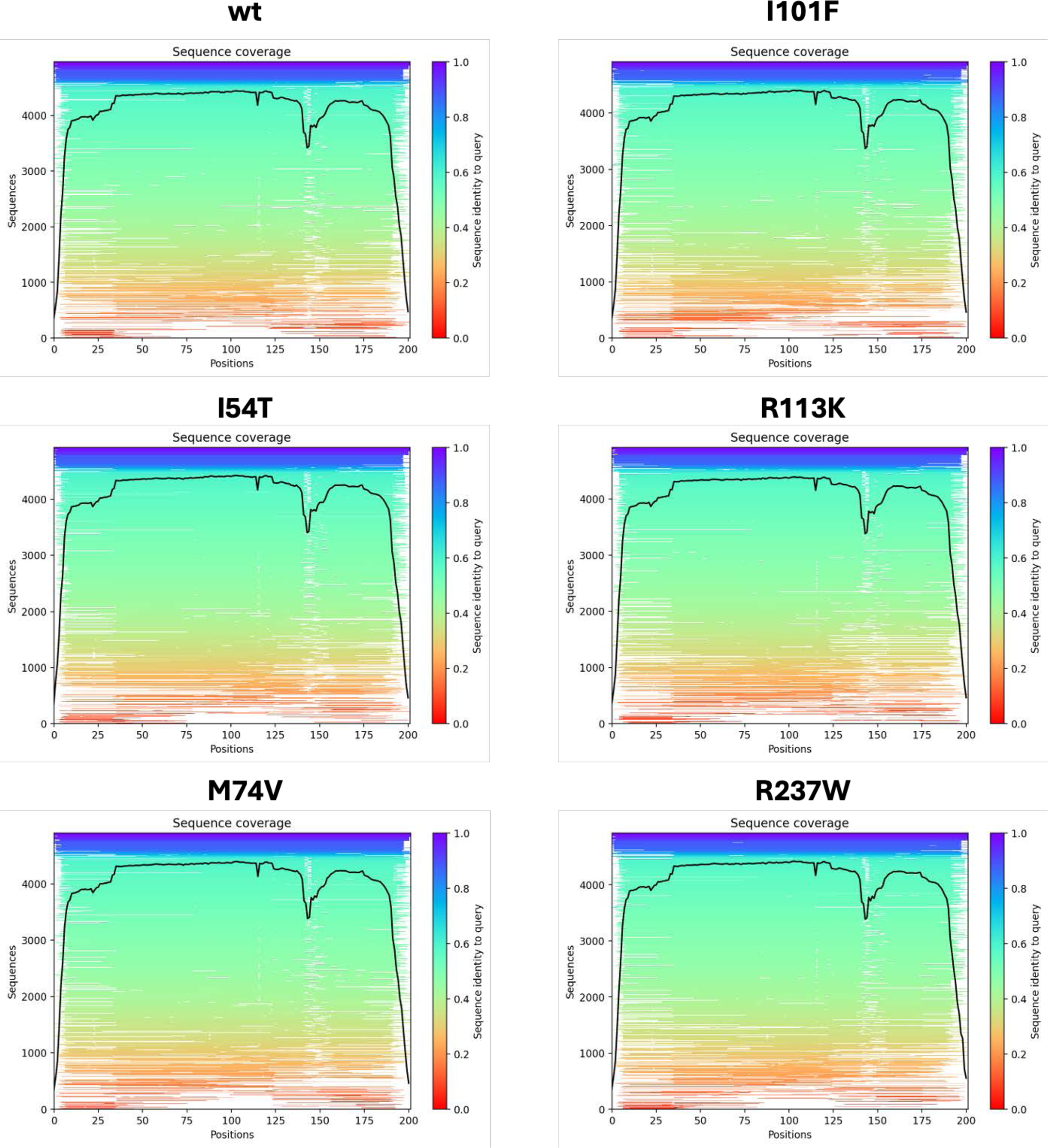
Sequence identity coverage of T-box domain and mutant structural predictions through Alphafold2.

**Supplementary Figure 3:**
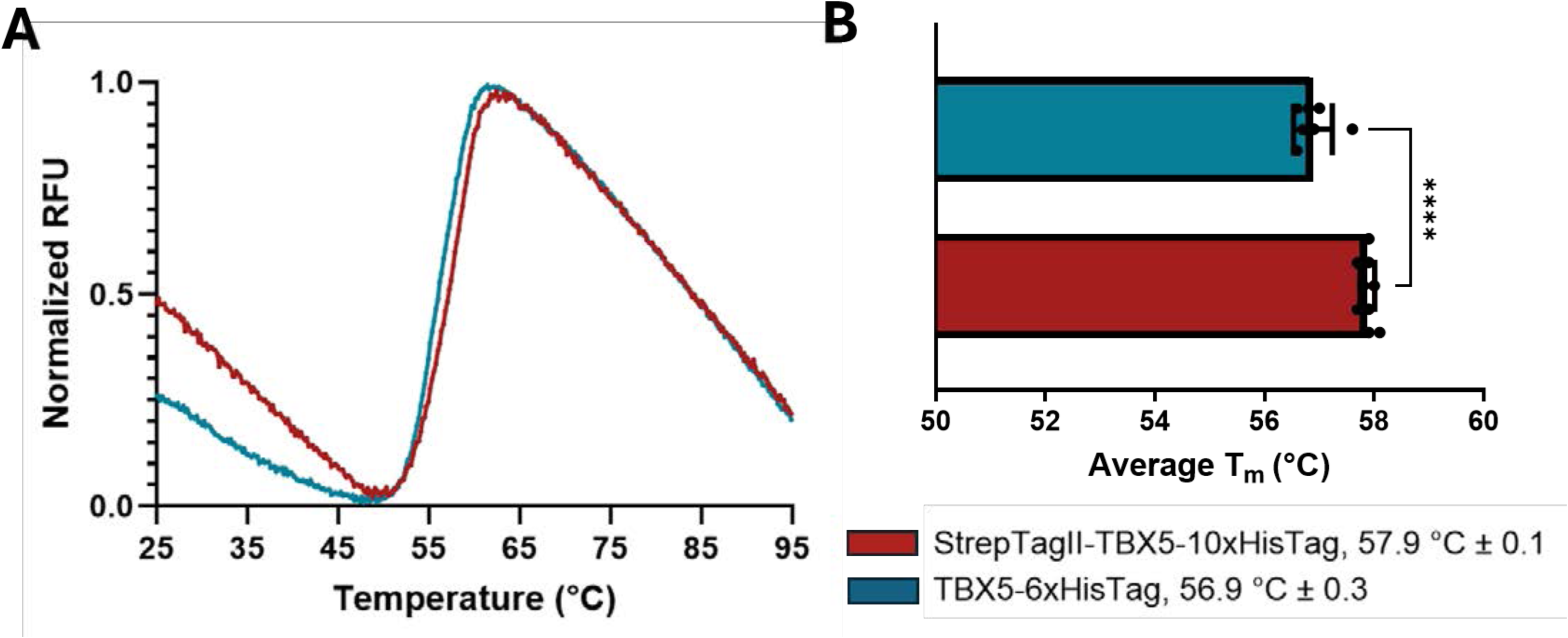
Wt T-box domain melting curves created through DSF. Melting curve assays were optimized using wt T-box domains with different purification tags. Color scheme: wt T-box domain with N-terminal strep tag and C-terminal 10x His-tag used in this work (red), wt T-box domain with a 6x His-Tag (blue).

**Supplementary Figure 4:**
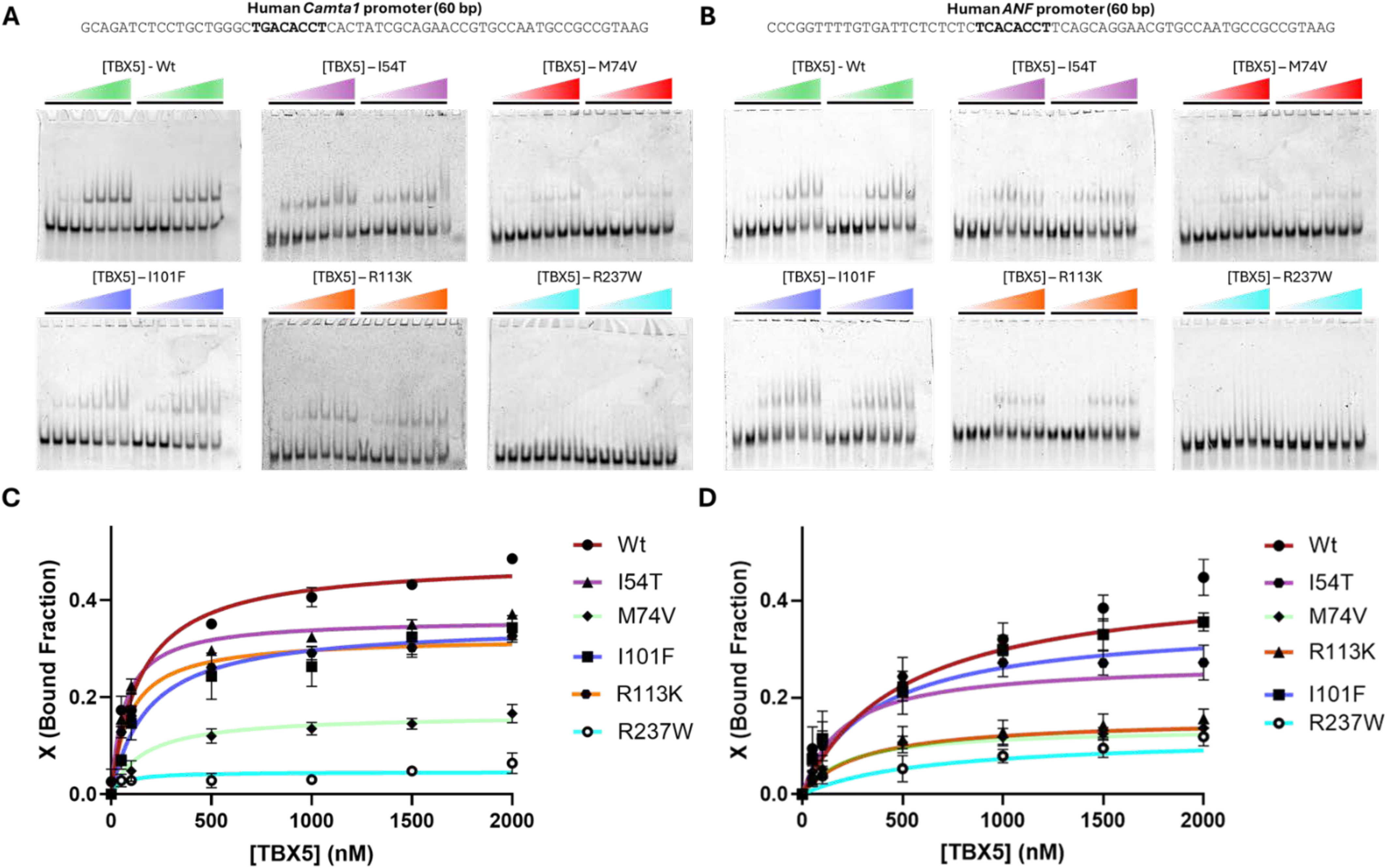
Representative EMSA gels used in this work A representative EMSA for wt T-box domain and mutants to evaluate binding to **A**) *Nppa* and **B**) *Camta1* to generate binding curves for Figure 4. Merged binding curves for **C**) *Nppa* and **D**) *Camta1* were generated to compare binding for all five missense mutants. Color scheme: Wt T-box domain (green), I54T (purple), M74V (red), I101F (dark blue), R113K (orange), and R237W (light blue/cyan). Sequences used for fluorescent probes are at the top of each figure with the wt TBX5 binding motif in bold.

**Supplementary Table 1:**
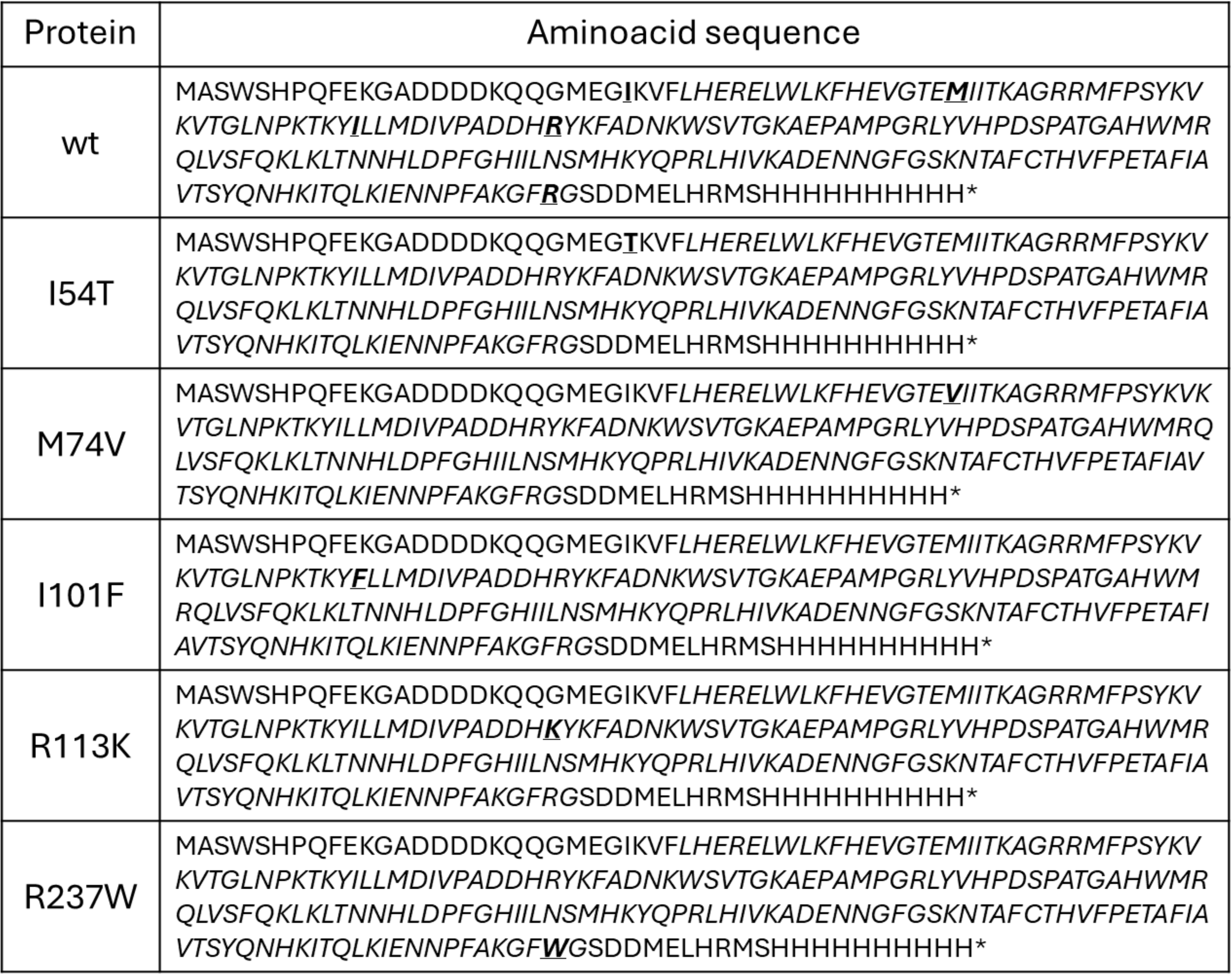
Amino acid sequence for wt and mutant T-box domain used in this work. Missense mutations are marked as underline amino acids.

**Supplementary Table 2:**
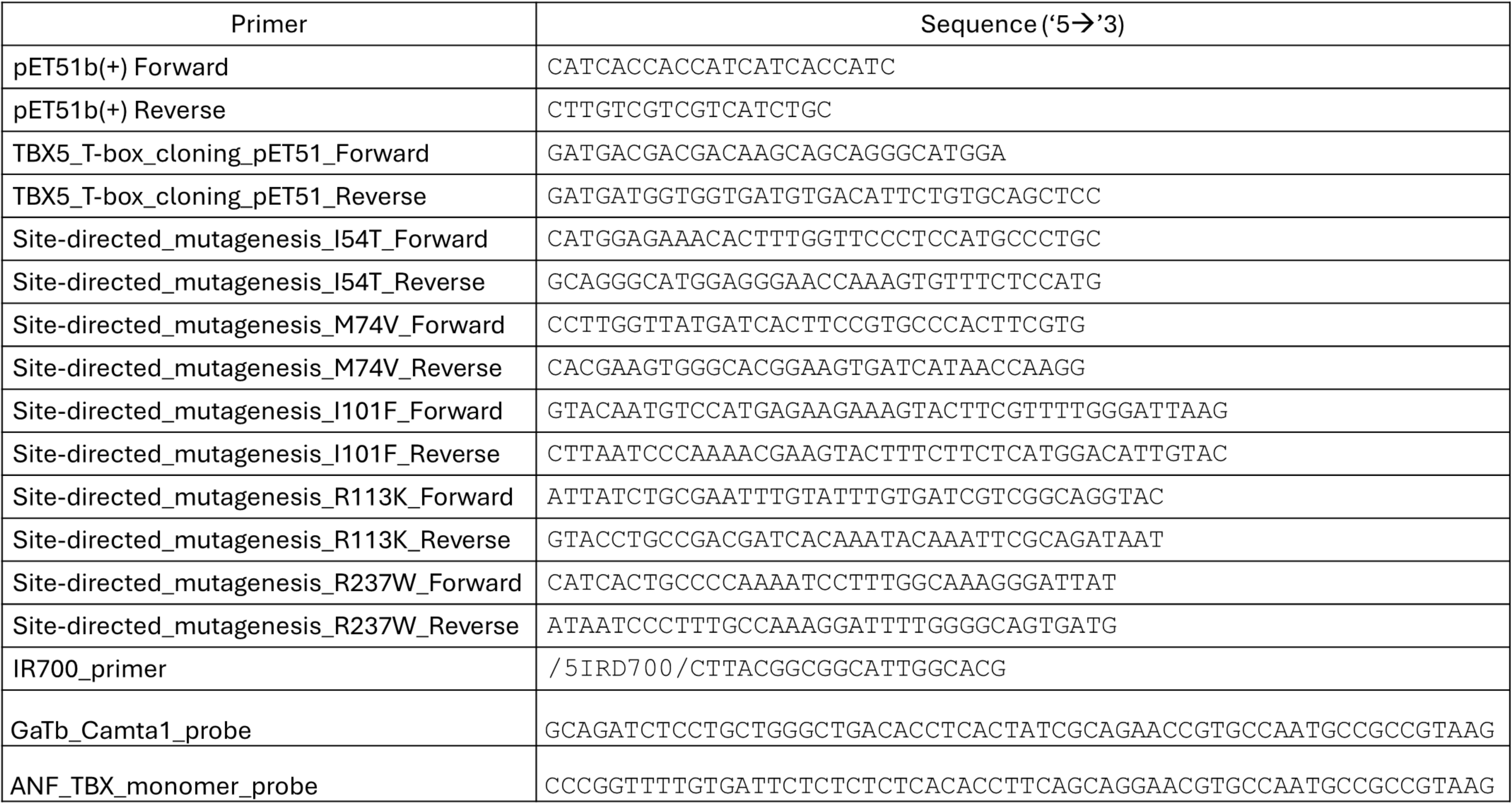
List of oligos used in this work.

**Supplementary Table 3:** Pathogenicity predictions through MutPred2. Gain of function and features are highlighted as green, while losses are highlighted as red.

| Mutation | MutPred2 Score (0,1) | Predicted pathogenic mechanism (p-value ≤ 0.05) | | |
| --- | --- | --- | --- | --- |
|  |  | Mechanism | Probability | p-value |
| I54T | 0.715 | Altered Metal binding | 0.17 | 0.04 |
|  |  | Altered Stability | 0.11 | 0.04 |
| M74V | 0.924 | Gain of Allosteric site at T72 | 0.28 | 3.7e-03 |
|  |  | Altered Metal binding | 0.24 | 0.02 |
|  |  | Altered DNA binding | 0.22 | 0.02 |
|  |  | Gain of Methylation at K78 | 0.17 | 9.7e-03 |
|  |  | Loss of Catalytic site at E73 | 0.12 | 0.03 |
| I101F | 0.789 | Gain of Allosteric site at D105 | 0.29 | 2.2e-03 |
|  |  | Altered Metal binding | 0.26 | 0.02 |
|  |  | Loss of Relative solvent accessibility | 0.26 | 0.03 |
|  |  | Loss of Acetylation at K99 | 0.25 | 0.01 |
|  |  | Loss of Catalytic site at D105 | 0.11 | 0.03 |
|  |  | Loss of Methylation at K99 | 0.11 | 0.03 |
| R113K | 0.800 | Gain of Relative solvent accessibility | 0.29 | 0.01 |
|  |  | Altered Metal Binding | 0.27 | 7.6e-03 |
|  |  | Gain of Acetylation at R113 | 0.27 | 6.3e-03 |
|  |  | Altered Ordered interface | 0.24 | 0.04 |
|  |  | Altered DNA binding | 0.19 | 0.03 |
|  |  | Gain of Methylation at K115 | 0.10 | 0.04 |
| R237W | 0.874 | Loss of Intrinsic disorder | 0.41 | 0.02 |
|  |  | Altered DNA binding | 0.34 | 5.0e-04 |
|  |  | Gain of Allosteric site at R237 | 0.33 | 7.0e-04 |
|  |  | Altered Disordered interface | 0.28 | 0.03 |
|  |  | Loss of Helix | 0.28 | 0.03 |
|  |  | Loss of Acetylation at K234 | 0.28 | 5.6e-03 |
|  |  | Gain of Loop | 0.27 | 0.04 |
|  |  | Loss of Methylation at K234 | 0.18 | 9.4e-03 |
|  |  | Gain of Catalytic site at K234 | 0.13 | 0.03 |

